# NEXCISION: exact, validated, and scalable excision of genomic regions from phylogenomic NEXUS matrices

**DOI:** 10.64898/2026.07.26.740842

**Authors:** Rhys T. White

## Abstract

Coordinate-based exclusion of genomic regions is routine in microbial phylogenomics, yet the editing step often relies on *ad hoc* scripts or manual alignment manipulation. This simple but high-consequence task can alter the character matrix, retain unwanted signal, or leave invalid NEXUS dimensions. This study presents NEXCISION, a dependency-free Python command-line tool for exact removal of coordinate-labelled rows from transposed NEXUS matrices. NEXCISION uses 1-based inclusive intervals, preserves retained rows and surrounding NEXUS content, updates matrix dimensions, reports interval-specific removal counts, and can generate deterministic provenance reports with SHA-256 checksums. Correctness was evaluated using three genuine SPANDx-derived matrices, an independent oracle, 17 correctness and preservation cases, 26 malformed-input challenges, 1,000 synthetic matrices containing 304,000 site rows, nine property-based invariants across 1,800 examples, and 12 deliberately faulted implementations. All expected rows were removed, retained rows and state strings were preserved, malformed inputs failed safely, and all deliberate faults were detected. In 105 measured scalability runs, NEXCISION remained correct and deterministic across 21 configurations. Runtime scaled near-linearly from ∼100 MB to 5 GB, processing a 4.83 GiB matrix in ∼24.2 seconds on one pinned logical central processing unit. Up to 100,000 intervals added modest runtime, while peak memory increased by ∼4.85 GiB per GiB of input. NEXCISION makes a fragile bespoke editing step exact, testable, and provenance-rich for microbial phylogenomics.

**Graphical abstract:** 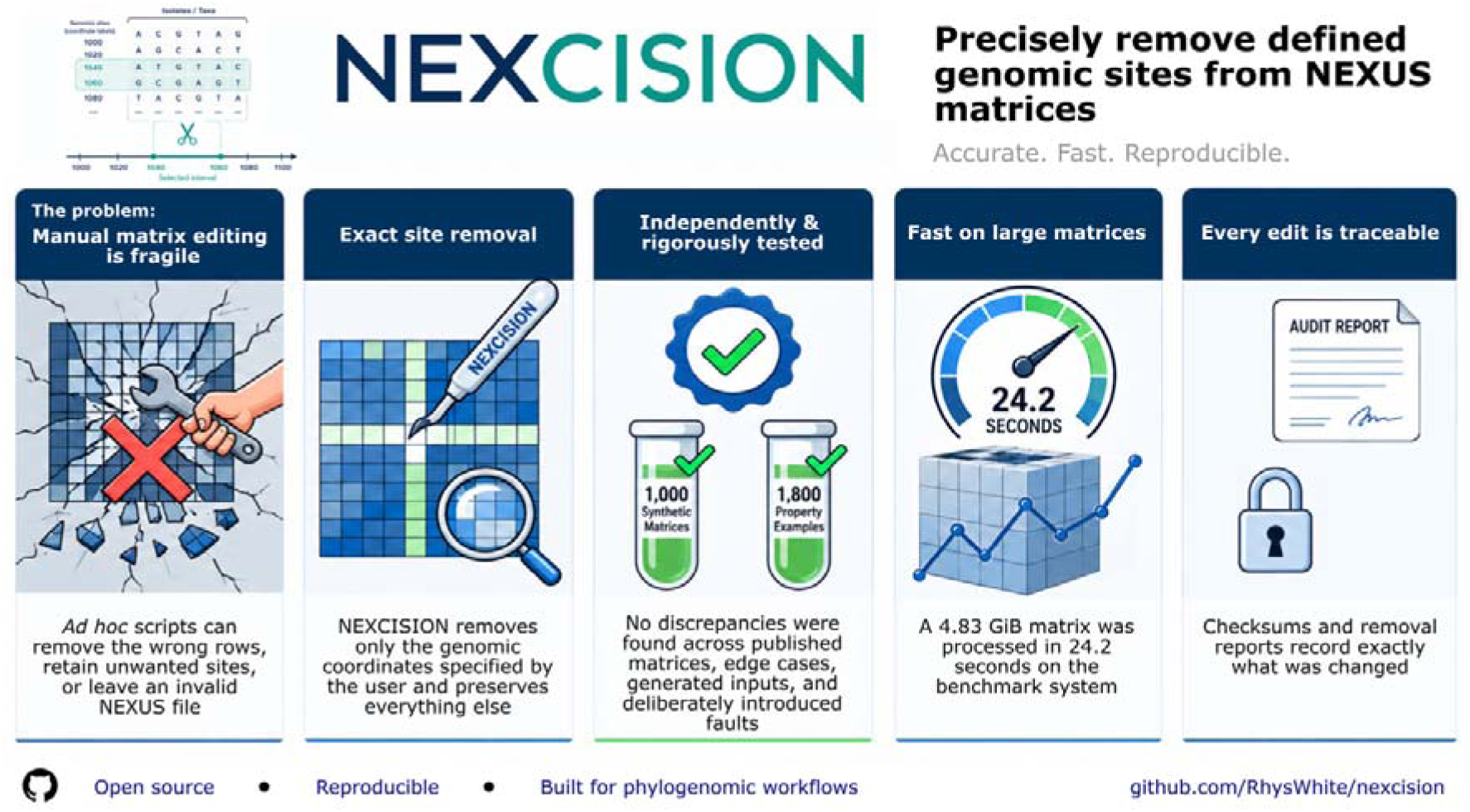

**Impact statement:** Removing defined genomic regions from a phylogenomic matrix seems simple, but errors can alter alignments, retain unwanted signal, or leave invalid NEXUS dimensions. NEXCISION replaces manual or bespoke editing with a focused command-line tool that removes only intended coordinate-labelled rows, preserves all other content, and records the operation. It is a trustworthy bridge between region detection and downstream phylogenetic inference for microbial genomic epidemiology, outbreak analysis, recombination masking, mobile-element studies, and workflows that excise reference-defined sites from transposed NEXUS matrices. NEXCISION was tested against genuine files, an independent oracle, 1,000 synthetic matrices, generated edge cases, and deliberately faulted implementations. It runs locally as a dependency-free, single-process Python utility. On a computer with 16 GB random-access memory (RAM), ∼2 GB matrices are practical; the largest tested matrix was 5.2 GB and required ∼25 GB RAM. NEXCISION is compact, transparent, and supported by evidence for correctness, failure safety, and scalability.

**Data summary:** No new biological sequence data were generated. Supporting software, validation assets, benchmarks, analysis code, figure sources, and reproducibility instructions are available from:

NEXCISION software repository: https://github.com/RhysWhite/nexcision

NEXCISION benchmarking repository: https://github.com/RhysWhite/nexcision-benchmarking

The evaluated software was NEXCISION v0.1.0 (MIT licence; Python >=3.10). The benchmarking repository includes the wheel checksum, deterministic data generator, manifests, independent oracle, compact results, summaries, and analysis scripts. The 1,000 synthetic matrices are reproducible from the version-controlled generator and manifest; the regenerated corpus has deterministic tree digest `9ed25d7e2dd4fee380f2f0f32ddcf653694715926905bd54179e7003e64f1c28`. Three SPANDx-derived matrices were used only to confirm compatibility with real matrix syntax; no biological inference was made. Their metadata and checksums are recorded in the benchmarking repository. All data, code, and protocols needed to reproduce the deterministic and performance results are provided in the article, supplementary material, or linked repositories. Software versions, computational environments, deterministic controls, and immutable provenance identifiers are summarized in Table S6.

## 6. Introduction

Reference-based single-nucleotide variant (SNV) analysis from whole-genome sequencing (WGS) is widely used in bacterial genomic epidemiology, especially for outbreak investigation and isolate-relatedness assessment (1–5). Pipelines such as SPANDx (6), SNVPhyl (1), and NASP (7) generate SNV alignments or matrices for phylogenetic reconstruction. Depending on the workflow, sites may be excluded if they fall within predicted regions of recombination, repetitive regions, or user-specified masks, including pre-identified phage or genomic-island regions (1, 8). Individual nucleotide calls may instead be masked for insufficient read depth, allele proportion, mapping quality, or other quality criteria (1, 6, 9, 10). Variants in paralogous loci may remain unresolved or be missed when short reads cannot be mapped to the reference genome unambiguously (6). Gubbins identifies substitution-dense regions consistent with recombination and generates a recombination-filtered phylogeny and alignment (11). However, when coordinate-defined exclusions are identified after a NEXUS matrix has been generated, applying them to that matrix requires an additional editing step (6, 12).

The NEXUS file format is a modular, extensible ASCII format for exchanging systematic data between programs (13). Its block-based structure can represent character matrices and trees, and link sequence alignments to masks, weights, comments, and other supplementary information, supporting exchange and archival of phylogenetic datasets (13, 14). Microbial genomics pipelines such as SPANDx use NEXUS to provide SNV matrices for phylogenetic analysis (6). In transposed NEXUS matrices, taxa are columns and characters are rows; in coordinate-labelled SNV matrices, each row can represent a genomic site defined by a reference sequence name and coordinate. This enables coordinate-based removal of sites within specified intervals, but reliable editing requires consistent label parsing, correct handling of inclusive boundaries and overlapping intervals, preservation of non-target content, and updating of the declared character count. Errors in any step can yield an incorrect or internally inconsistent phylogenetic matrix.

Although coordinate-based matrix editing is conceptually simple, its correctness can be tested against an exact expected result: every target row should be removed, every non-target row retained, and matrix structure preserved. Bacterial phylogenomics benchmarks have used simulated datasets with known variants to quantify variant-calling accuracy (15), and some have compared inferred trees with the phylogenies used to generate the simulated data to assess topological accuracy (7, 16). Property-based testing checks specified software properties against automatically generated test cases (17, 18), while mutation testing evaluates whether tests detect deliberately introduced faults (19).

This study introduces NEXCISION (https://github.com/RhysWhite/nexcision), a dependency-free command-line tool that removes reference-coordinate-labelled rows from transposed NEXUS SNV matrices. Using inclusive coordinates, NEXCISION resolves overlapping intervals without double-counting, removes only valid matching rows, preserves non-target content, updates the declared character count, and can generate provenance reports of intervals and removed sites. It was evaluated against exact expected outputs using fixed and random tests, property-based and mutation testing, and published SPANDx-derived matrices (20–22). Runtime and memory were benchmarked across increasing matrix and exclusion-set size. NEXCISION provides an auditable bridge between coordinate-based region identification and structurally consistent NEXUS matrices for downstream phylogenetic analysis.

## 7. Theory and implementation

### 7.1 Supported matrix model and coordinate semantics

NEXCISION v0.1.0 is a dependency-free Python command-line application requiring Python 3.10 or newer. It is designed primarily for a transposed NEXUS matrix block in which each complete matrix row represents a genomic site and the first token ends with a coordinate. By default, the terminal integer following an underscore is extracted using the regular expression `_(\d+)$`; users may supply an alternative expression containing exactly one capture group.

Region files are whitespace-delimited and contain start and end coordinates with an optional name. Coordinates are interpreted as 1-based and inclusive. Reversed interval boundaries are normalized automatically. NEXCISION assumes one shared reference-coordinate axis per matrix: it does not currently distinguish contigs or replicons that reuse the same coordinate values. That deliberately narrow scope is checked explicitly rather than inferred silently.

### 7.2 Filtering algorithm

Input intervals are normalized, sorted, and merged into a union of overlapping or directly adjacent spans (start positions are indexed). Each parsed matrix coordinate is then evaluated against the merged union by binary search, giving efficient membership testing without repeatedly scanning every supplied interval. The original, unmerged intervals are retained separately so that per-region removal counts can be reported in the order supplied by the user. A matrix row matching several overlapping regions is removed once but contributes independently to each corresponding regional count.

Rows outside the interval union are written without alteration and remain in their original order. Comments, blank lines, command capitalization, state delimiters and content outside the matrix block are preserved. For a transposed matrix, NEXCISION updates `nchar` only when the declared value equals the original number of matrix rows; `ntax` is preserved. If the relevant dimension is inconsistent with the observed row count, the software warns rather than guessing.

### 7.3 Command-line interface, outputs and safety model

A typical invocation is:

nexcise input.nex regions.tsv \

--output filtered.nex \

--counts removed_counts_per_region.tsv \

--report nexcision_report.json

The principal output is a filtered NEXUS file. A tab-delimited counts file reports the number of removed rows associated with every supplied interval. An optional deterministic JSON report records parameters, observed dimensions, warnings, removal totals and SHA-256 checksums for inputs and outputs. Report fields can be inspected programmatically to enforce workflow-specific acceptance criteria, such as an expected removal count, the absence of warnings, or agreement with recorded input and output checksums. Example shell and Snakemake integrations that parse report fields, enforce analysis-specific acceptance criteria, and verify recorded SHA-256 checksums are provided in the /`docs/workflow-integration.md` and `/examples/snakemake/` available in the NEXCISION GitHub repository (https://github.com/RhysWhite/nexcision/). The supported input structure, coordinate-based excision, and resulting matrix and audit outputs are summarized in Fig. 1. Existing outputs are not overwritten unless `--force` is supplied.

**Figure 1.**
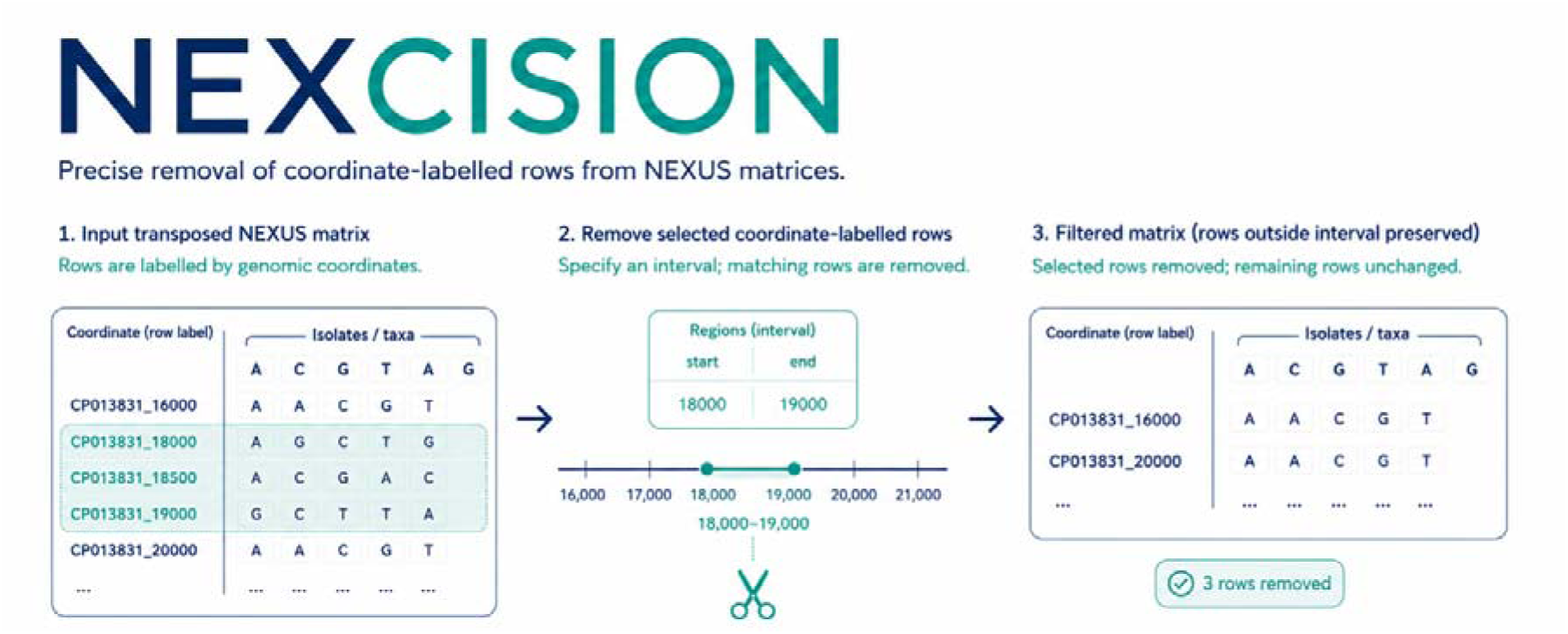
NEXCISION workflow. A transposed NEXUS matrix contains coordinate-labelled site rows. User-defined 1-based inclusive intervals select matching rows for removal; all rows outside the interval are retained unchanged. The matrix dimension is updated and optional count and provenance files record the operation.

Safety checks reject missing or multiple matrix blocks, unterminated matrices, invalid UTF- 8, malformed region coordinates, unsafe output paths and unparseable matrix rows by default. Output files are staged before commit so that a failed multi-output operation does not leave a partial result. Filtering that would remove every matrix row is rejected unless `--allow-empty` is supplied explicitly. These behaviors follow the broader principles of usable command-line bioinformatics and reproducible computational research (23, 24).

### 7.4 Validation framework

Validation was structured as complementary evidence layers rather than a single test set (Fig. 2). First, three genuine SPANDx matrices established compatibility with real outputs: 47 taxa and 266 site rows generated using SPANDx v4.0.4 (22); 237 taxa and 4,649 site rows generated using SPANDx v3.2 (20); and 396 taxa and 10,481 site rows generated using SPANDx v4.0.4 (21). Exact-site and short-interval tests were independently reconstructed for each matrix (Table S1).

**Figure 2.**
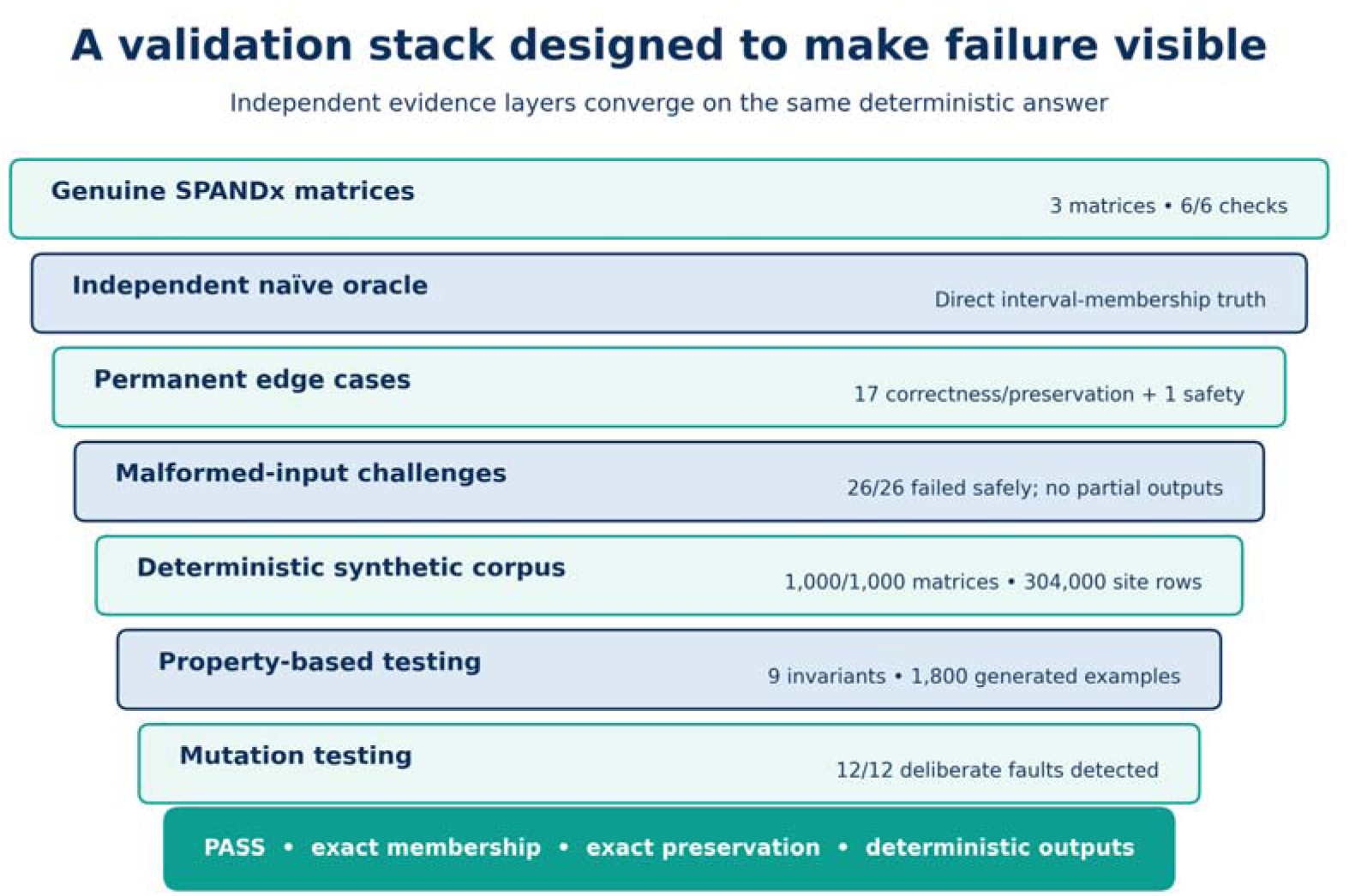
Validation framework and outcomes. Seven complementary evidence layers targeted distinct failure modes in coordinate-based matrix editing. All prespecified correctness, preservation, failure-safety, and mutation-detection criteria were met.

Second, an independent naïve oracle calculated expected membership using the direct rule start <= coordinate <= end and imported no NEXCISION code. Seventeen permanent cases tested inclusive boundaries, sites immediately outside an interval, overlapping, duplicate, adjacent, reversed, unsorted and no-match regions, row order, comments, blank lines, state spacing, command capitalization, and inline matrix terminators. A separate all-sites case tested empty-output protection. Twenty-six malformed-input cases tested clean failure and transactional output safety (Table S2).

Third, a deterministic corpus of 1,000 synthetic transposed NEXUS matrices was generated across ten scenario families. Matrices contained 2–256 taxa, 20–1,000 site rows, one to four regions and 0–80% row removal, with ascending, descending or shuffled row order. Exact results were compared with the independent oracle; the complete corpus design and scenario- family outcomes are provided in Table S3. Fourth, Hypothesis v6.158.0 (18) generated 1,800 additional examples across nine invariants, including membership, preservation, dimensions, idempotence, interval-order invariance, overlap equivalence, all-site safety, and custom coordinate patterns. Finally, 12 plausible source mutations were introduced into temporary implementations, including exclusive interval boundaries, wrong-dimension updates, incomplete interval merging, incorrect coordinate extraction, row reordering, lost terminators, disabled empty-output protection, and incorrect regional counts. Mutation testing asks whether a test suite can detect deliberately introduced faults (19). The complete property-based invariants and deliberate source mutations are detailed in Table S4.

Because matrix editing is deterministic, all correctness criteria were prespecified as exact. No power calculation, statistical hypothesis test, or blinding was applicable. Synthetic generation and benchmark run order were deterministic or explicitly seeded.

### 7.5 Scalability benchmarking

Scalability was evaluated using the frozen NEXCISION v0.1.0 wheel (https://github.com/RhysWhite/nexcision-benchmarking/blob/959859aa444784c9cedac5088520b322ded49567/software/nexcision-0.1.0-py3-none-any.whl) on an exclusively allocated Linux node containing two AMD EPYC 9374F sockets, with the NEXCISION process pinned to one logical central processing unit (CPU). Inputs and outputs were stored on node-local XFS storage. The benchmark used one unreported warm-up followed by five measured repetitions for each of 21 configurations, giving 105 measured executions. Measured-run order was randomized with a fixed seed. Page cache was not dropped; the results therefore represent controlled warm-cache application performance rather than cold network-storage performance.

Six configurations varied input size from approximately 10 MB to 5 GB. Three ∼500 MB matrices separated the effect of matrix shape by contrasting many short rows, a balanced matrix and fewer very wide rows. Six configurations varied the exclusion list from 1 to 100,000 intervals on an ∼1 GiB matrix. Four configurations compared disjoint, adjacent, overlapping and nested interval geometry at 10,000 submitted intervals. Two configurations measured the overhead of the provenance report. Wall time, user and system CPU time, peak resident memory, throughput, output size, dimensions, and checksums were recorded. Median and interquartile range were used descriptively across five repetitions. Linear models were fitted to configuration medians for input-size scaling. Complete configuration-level designs and outcomes are provided in Table S5.

## 8. Results

### 8.1 Exact excision and preservation across validation layers

All six checks on genuine SPANDx matrices removed exactly the intended rows. Across these checks, retained row labels, state strings, and order were unchanged; `ntax` remained constant; `nchar` was reduced correctly; and content following the matrix block was preserved.

All 17 permanent correctness and preservation cases passed, the all-sites safety case behaved as specified, and all 26 malformed-input cases failed cleanly without modifying inputs or leaving unintended outputs. In the 1,000-case synthetic corpus, NEXCISION evaluated 304,000 site rows, removing 113,740 and retaining 190,260. There were no disagreements with the independent oracle, altered retained rows, dimension errors, regional-count errors, checksum discrepancies or repeat-run differences. All nine property-based invariants passed across 1,800 generated examples, and all 12 deliberate source faults were detected, giving a mutation score of 100% (Table 1; Fig. 2).

**Table 1.** Formal correctness and failure-safety validation of NEXCISION v0.1.0.

| Validation component | Result |
| --- | --- |
| Genuine SPANDx matrices | 3 files; 6/6 checks passed |
| Unit tests | 26/26 passed |
| Fixed correctness and preservation cases | 17/17 passed |
| All-sites safety case | 1/1 passed |
| Malformed-input and output-safety cases | 26/26 passed |
| Deterministic synthetic matrices | 1,000/1,000 passed |
| Property-based testing | 9/9 invariants; 1,800 generated examples |
| Deliberate source mutants | 12/12 detected; 100% mutation score |

### 8.2 Runtime and memory scaling

All 105 formally measured executions were correct, and each of the 21 configurations produced deterministic output checksums across five randomized repetitions. Excluding the ∼10 MB configuration, where sub-second process-startup noise inflated variability, the median runtime coefficient of variation across configurations was 1.08%. Normalized runtime showed no material drift with randomized run order.

Runtime increased near-linearly with input size. Median elapsed time was 0.52 seconds for ∼100 MB, 4.51 seconds for ∼1 GB, 8.86 seconds for ∼2 GB, and 24.24 seconds for a 4.83 GiB matrix. A linear model fitted to configuration medians explained 99.86% of runtime variation; the corresponding log-log exponent for inputs of at least ∼100 MB was 0.978 (Fig. 3A).

**Figure 3.**
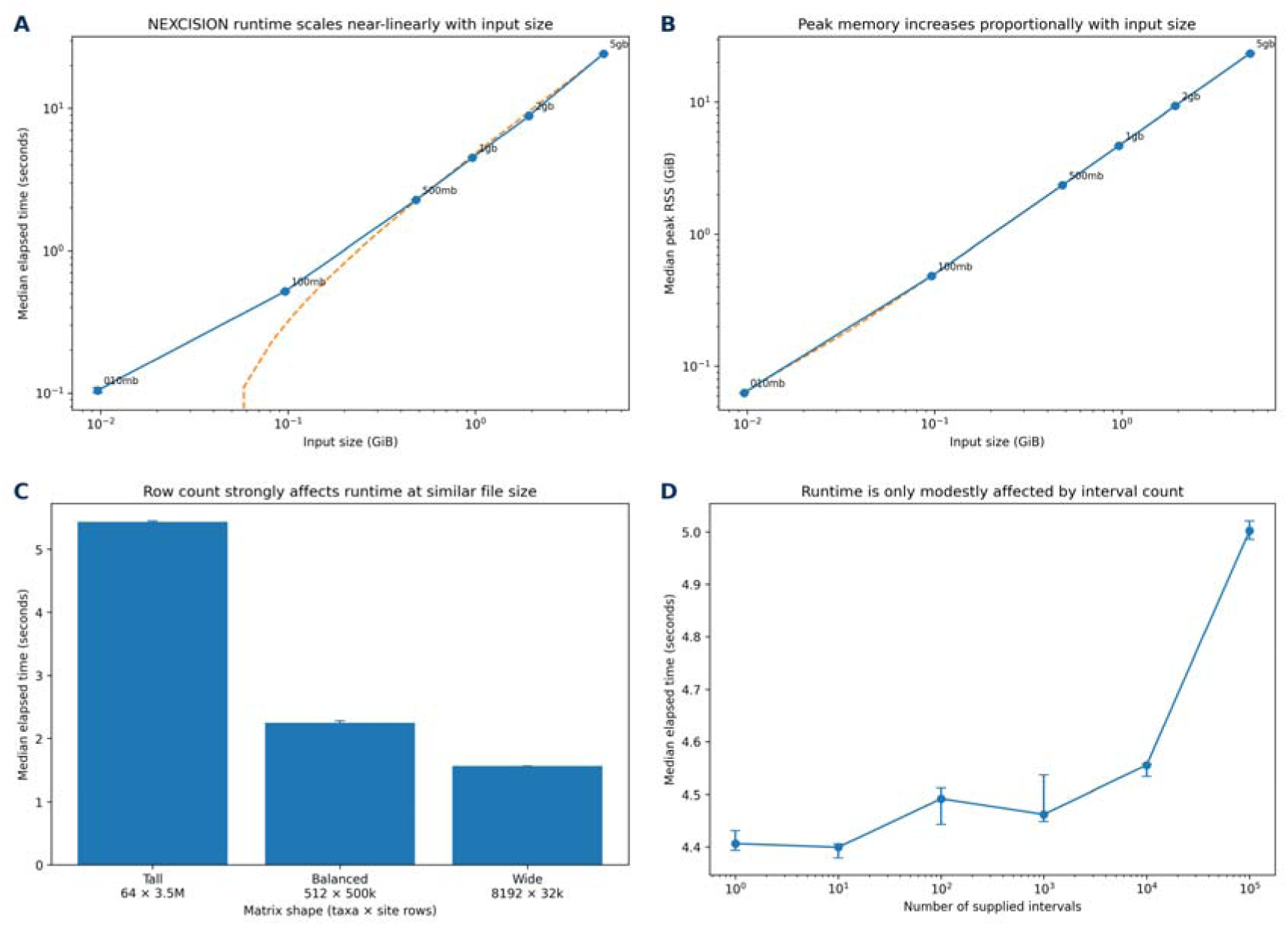
Scalability and resource use of NEXCISION. (**A**) Median elapsed time as a function of input-matrix size. (**B**) Median peak resident memory as a function of input size. Dashed lines in A and B show linear fits to configuration medians; both axes are logarithmic. (**C**) Effect of matrix shape at broadly similar file size. (**D**) Effect of exclusion-list size on elapsed time for a fixed approximately 1 GiB matrix. Points or bars represent medians of five randomized measured executions; error bars show interquartile ranges. Benchmarks used NEXCISION v0.1.0 on one pinned logical CPU of an exclusively allocated AMD EPYC 9374F Linux node with node-local XFS storage.

Peak resident memory also increased linearly with input size. The fitted slope was ∼4.85 GiB of peak Resident Set Size (RSS) per GiB of input, with an R² effectively equal to 1.0. The 4.83 GiB matrix required ∼23.4 GiB peak RSS (Fig. 3B). Thus, the current implementation was fast at multi-gigabyte scale but was not memory-light.

### 8.3 Effects of matrix shape, interval complexity, and provenance

Matrix shape affected runtime independently of file size. Among three matrices of ∼500 MB, the tall matrix containing 3.5 million site rows required 5.44 seconds, compared with 2.25 seconds for the balanced matrix and 1.56 seconds for the wide matrix containing 32,000 site rows. The tall matrix was therefore 3.48-fold slower than the wide matrix, indicating substantial per-row parsing and coordinate-evaluation overhead (Fig. 3C).

Increasing the exclusion list from one to 100,000 intervals increased median runtime by 13.5% and peak memory by 0.8% (Fig. 3D). At 10,000 submitted intervals, disjoint, adjacent, overlapping and nested geometries differed in median runtime by 2.8%, indicating little practical penalty from interval arrangement after preprocessing. Enabling the deterministic JSON provenance report increased median runtime from 4.43 to 5.48 seconds for the ∼1 GiB matrix, a 23.6% overhead, while peak memory was effectively unchanged. Selected scalability outcomes are summarized in Table 2.

**Table 2.** Selected scalability outcomes from the formal benchmark.

| **Benchmark outcome** | **Observed result** |
| --- | --- |
| Correctness and determinism | 105/105 measured runs correct; 21/21 configurations deterministic |
| Runtime scaling | Near-linear; $R^2=0.9986$; log-log exponent=0.978 for $\geq 100$ MB |
| Largest tested input | 4.83 GiB processed in 24.24 s |
| Peak-memory scaling | $\sim 4.85$ GiB RSS per GiB input |
| Matrix-shape effect | Tall matrix 3.48-fold slower than wide matrix at similar size |
| Interval-count effect | 1 to 100,000 intervals increased runtime by 13.5% |
| Interval-geometry effect | 2.8% spread across disjoint, adjacent, overlapping, and nested |
| Provenance-report overhead | 23.6% additional runtime; peak memory unchanged |

## 9. Discussion

NEXCISION (https://github.com/RhysWhite/nexcision) addresses a narrow operation with disproportionate analytical consequences. Established tools can identify regions for exclusion and infer phylogenetic trees, but applying those coordinates to an existing NEXUS matrix may still depend on one-off scripts. Any errors introduced at this stage are carried directly into the matrix used for phylogenetic analysis. NEXCISION makes the operation explicit through user-defined intervals, inclusive coordinate semantics, exact row membership, preservation of retained content, dimension-aware output, and an optional machine-readable provenance record.

Genomic filtering can be performed at several stages of an analysis, but filters applied before and after variant calling operate on different data types and should not be treated as equivalent (25). SNVPhyl, for example, accepts user-defined and internally generated repeat masks, but maps reads to the unmodified reference before SNVs at masked coordinates are excluded during downstream alignment and phylogeny generation (1). Other tools act later in the workflow: maskrc-svg applies ClonalFrameML (26) or Gubbins-derived recombinant- region masks to an existing multiple-sequence alignment (https://github.com/kwongj/maskrc-svg/tree/master). By contrast, altering or truncating the mapping reference changes the sequence against which reads are aligned and may create artificial boundaries. Reference divergence, incomplete reference sequence, repeats, and structural differences can affect read placement and SNV-calling accuracy, while reads spanning truncated reference boundaries may align only partially or fail alignment filters (27, 28). NEXCISION occupies a distinct post-mapping role: it applies an already defined coordinate mask to an existing matrix without altering upstream read mapping or variant calling.

The strongest result is not that NEXCISION produced the expected output on several example files. Independent and adversarial validation found no discrepancy across published matrices (20–22), permanent edge cases, malformed inputs, 304,000 synthetic site rows, 1,800 generated property examples, and 12 deliberately introduced faults. Together, these tests make the correctness claim falsifiable. No finite test suite can prove the absence of every possible defect, but agreement among an independent oracle, fixed edge cases, generated inputs, malformed-input challenges, and deliberately broken implementations provides substantially stronger evidence than a small set of successful examples. Using multiple validation layers is especially important because comparative benchmarking of microbial SNV pipelines has shown that apparent performance can vary across datasets, reference genomes, and internal filtering strategies, and that results from a single benchmark may not generalize (29). The validation framework also offers a practical model for validating small scientific utilities: define a narrow scope, specify exact invariants, separate the oracle from the implementation, and demonstrate that the tests fail when the software is intentionally broken.

NEXCISION v0.1.0 was computationally practical for large microbial matrices on a single logical CPU. Runtime increased near-linearly with input size, and increasing the exclusion list from one to 100,000 intervals added only modest runtime. Matrix shape nevertheless affected performance independently of file size: matrices containing many short site rows required more time than similarly sized matrices containing fewer, wider rows, consistent with per-row parsing and coordinate-evaluation overhead. Provenance-report generation imposed a moderate runtime cost but no measurable memory penalty.

Peak memory was the main limitation. NEXCISION v0.1.0 reads the complete UTF-8 file and constructs filtered content in memory, producing a ∼4.85-fold peak-RSS multiplier relative to input size in the tested format. The 4.83 GiB matrix therefore required ∼23.4 GiB peak RSS, making available memory rather than runtime the likely constraint on ordinary workstations and for still larger inputs. A streaming implementation and lower-copy output staging are priorities for a future release.

The supported input model is deliberately narrower than the complete NEXUS specification. NEXCISION expects complete coordinate-labelled rows in a single matrix block and, by default, a terminal integer coordinate. It does not currently distinguish multiple contigs or replicons that reuse the same numerical coordinates. Future collaborations and development could extend NEXCISION to commonly used variant representations, including Snippy (https://github.com/tseemann/snippy) generated multisample Variant Call Format (VCF) file and core-genome alignment outputs. Such support would require contig-aware interval definitions and coordinated filtering across linked output formats.

Masking is not an analytically neutral operation. Benchmarking of SNVPhyl and *Mycobacterium tuberculosis* workflows demonstrates a recurring trade-off: excluding broadly defined problematic regions can reduce false-positive calls and improve precision or specificity, but may also increase false negatives and reduce recall or sensitivity, including through the removal of positions that are otherwise accurately callable (1, 30). More generally, filtering choices and thresholds can substantially affect the data retained and the conclusions drawn from downstream analyses (25, 31). The appropriate mask therefore depends on the organism, sequencing data, reference genome, mapping workflow, and analytical objective. NEXCISION does not determine which regions should be excluded; it ensures that a supplied coordinate mask is applied exactly, reproducibly, and transparently. The present study validates that editing operation, not the biological validity of the supplied intervals or the downstream phylogenetic consequences of their removal.

Within this defined scope, NEXCISION replaces a fragile manual step with a reproducible and auditable workflow. Potential applications include excision of recombination blocks, prophages, mobile-element insertions, duplicated regions, low-confidence mapping intervals, and any reference-defined mask that must be applied after generation of a transposed NEXUS matrix. The software is small enough to audit, has no runtime dependencies, and can be incorporated into shell scripts or workflow managers. Machine-readable removal counts, warnings, dimensions, and checksums allow users to define analysis-specific validation rules; worked integration examples are provided in the NEXCISION GitHub repository (https://github.com/RhysWhite/nexcision). NEXCISION reduces the risk that a simple file-editing error undermines an otherwise rigorous phylogenomic analysis.

## 11. Author statements

### 11.1 Author contributions

RTW: conceptualization, software, methodology, validation, formal analysis, investigation, data curation, visualization, writing (original draft, writing), review, and editing, project administration.

### 11.4 Ethical approval

Not applicable. This study evaluated software using synthetic matrices and de-identified phylogenomic matrix outputs. It involved no experimental work with humans or animals and no identifiable personal data.

### 11.5 Consent for publication

Not applicable. The article contains no identifiable personal information.

## Acknowledgements

The author acknowledges Genomics Aotearoa and PHF Science for supporting the development and benchmarking of NEXCISION. The author also thanks colleagues working in genomic epidemiology and public health bioinformatics whose practical need to remove defined genomic regions from single-nucleotide variant matrices motivated this resource.

## 12. Abbreviations

ASCII: American Standard Code for Information Interchange
CPU: central processing unit
GB: gigabyte
GiB: gibibyte
MB: megabyte
JSON: JavaScript Object Notation
RAM: random-access memory
RSS: Resident Set Size
SNV: single-nucleotide variant
UTF-8: Unicode Transformation Format–8-bit
VCF: Variant Call Format
WGS: whole-genome sequencing

## 13. Declaration of generative AI and AI-assisted technologies in the manuscript preparation process

During the preparation of this work, the author used ChatGPT (OpenAI) to assist with minor copy editing and formatting of author-generated text, and with code debugging. All analytical decisions, code implementation, results, interpretations, and conclusions were developed, checked, and approved by the author. The author takes full responsibility for the content of the manuscript.

